# A reevaluation of the relationship between EGL-43 (EVI1/MECOM) and LIN-12 (Notch) during *C. elegans* anchor cell invasion

**DOI:** 10.1101/2022.10.04.510909

**Authors:** Michael A. Q. Martinez, Angelina A. Mullarkey, Callista Yee, Chris Z. Zhao, Wan Zhang, Kang Shen, David Q. Matus

**Affiliations:** Department of Biochemistry and Cell Biology, Stony Brook University, Stony Brook, NY 11794, USA; Howard Hughes Medical Institute, Department of Biology, Stanford University, Stanford, CA 94305, USA

**Keywords:** *C. elegans*, AC Invasion, EGL-43, LIN-12, AID, DHB

## Abstract

Development of the *C. elegans* reproductive tract is orchestrated by the anchor cell (AC). Among other things, this occurs through a cell invasion event that connects the uterine and vulval tissue. Several key transcription factors regulate AC invasion, such as EGL-43, HLH-2, and NHR-67. Specifically, these transcription factors function together to maintain the post-mitotic state of the AC, a requirement for AC invasion. EGL-43 is the *C. elegans* homolog of the human EVI1/MECOM proto-oncogene, and recently, a mechanistic connection has been made between its loss and AC cell-cycle entry. The current model states that EGL-43 represses LIN-12 (Notch) expression to prevent AC proliferation, suggesting that Notch signaling is mitogenic in the absence of EGL-43. To reevaluate the relationship between EGL-43 and LIN-12, we designed and implemented a heterologous co-expression system called AIDHB that combines the auxin-inducible degron (AID) system of plants with a live cell-cycle sensor based on human DNA helicase B (DHB). After validating the AIDHB approach using AID-tagged GFP, we sought to test this approach using AID-tagged alleles of *egl-43* and *lin-12*. Auxin-inducible degradation of either EGL-43 or LIN-12 resulted in the expected AC phenotypes. Lastly, we seized the opportunity to pair AIDHB with RNAi to co-deplete LIN-12 and EGL-43, respectively. This combined approach revealed that LIN-12 is not required for AC proliferation following loss of EGL-43, which contrasts with a double RNAi experiment directed against these same targets. The addition of AIDHB to the *C. elegans* transgenic toolkit should facilitate functional *in vivo* imaging of cell-cycle associated phenomena.

## Introduction

Cell invasion through basement membrane (BM) is essential for animal development, tissue inflammation, and cancer metastasis. During *C. elegans* larval development, a specialized uterine cell, the anchor cell (AC), breaches BM to contact the underlying vulval epithelium. This developmental event initiates the attachment of the uterus to the vulva, which later forms the reproductive tract of the animal. Several laboratories, including ours, have taken advantage of the animal’s simple anatomy, transparent body, and genetic amenability to characterize molecular and cellular features of *C. elegans* AC invasion. Collectively, this has yielded important insights into the regulation of BM invasion *in vivo* (Sherwood and Plastino, 2018).

One requirement for AC invasion is the maintenance of the post-mitotic state (Matus et al., 2015), which is executed by a network of conserved transcription factors that includes EGL-43 (EVI1/MECOM), HLH-2 (E/Daughterless), and NHR-67 (TLX/Tailless) (Deng et al., 2020; Medwig-Kinney et al., 2020). Together these three transcription factors form a coherent (type I) feed-forward loop with positive feedback (Medwig-Kinney et al., 2020). Loss of either EGL-43, HLH-2, or NHR-67 results in AC proliferation with defective BM invasion. Until recently, the mechanism connecting the loss of these transcription factors with AC proliferation was poorly understood. New research has revealed that EGL-43 maintains the post-mitotic state of the AC by repressing LIN-12 (Notch) expression (Deng et al., 2020), suggesting that Notch signaling promotes AC proliferation.

To reevaluate the relationship between EGL-43 and LIN-12 during AC invasion, we generated a heterologous co-expression system that allows conditional degradation of target proteins and visualization of cell-cycle state (Fig. 1A). Targeted protein degradation is triggered by the plant-derived auxin-inducible degron (AID) system (Nishimura et al., 2009), and the cell cycle is monitored using a biosensor based on human DNA helicase B (DHB) (Hahn et al., 2009; Martinez and Matus, 2022; Spencer et al., 2013). We tested the co-expression system, referred to as AIDHB, by degrading GFP as well as endogenous EGL-43 and LIN-12. We show that it is robust, as it strongly degrades GFP without causing AC cell-cycle defects and produces highly penetrant AC phenotypes associated with the loss of either EGL-43 or LIN-12. Finally, we combined AIDHB with RNAi to simultaneously deplete LIN-12 and EGL-43. Though we confirm that EGL-43 represses the endogenous expression of LIN-12 (Notch) during AC invasion, our results imply that LIN-12 is not required for AC proliferation.

**Figure 1.**
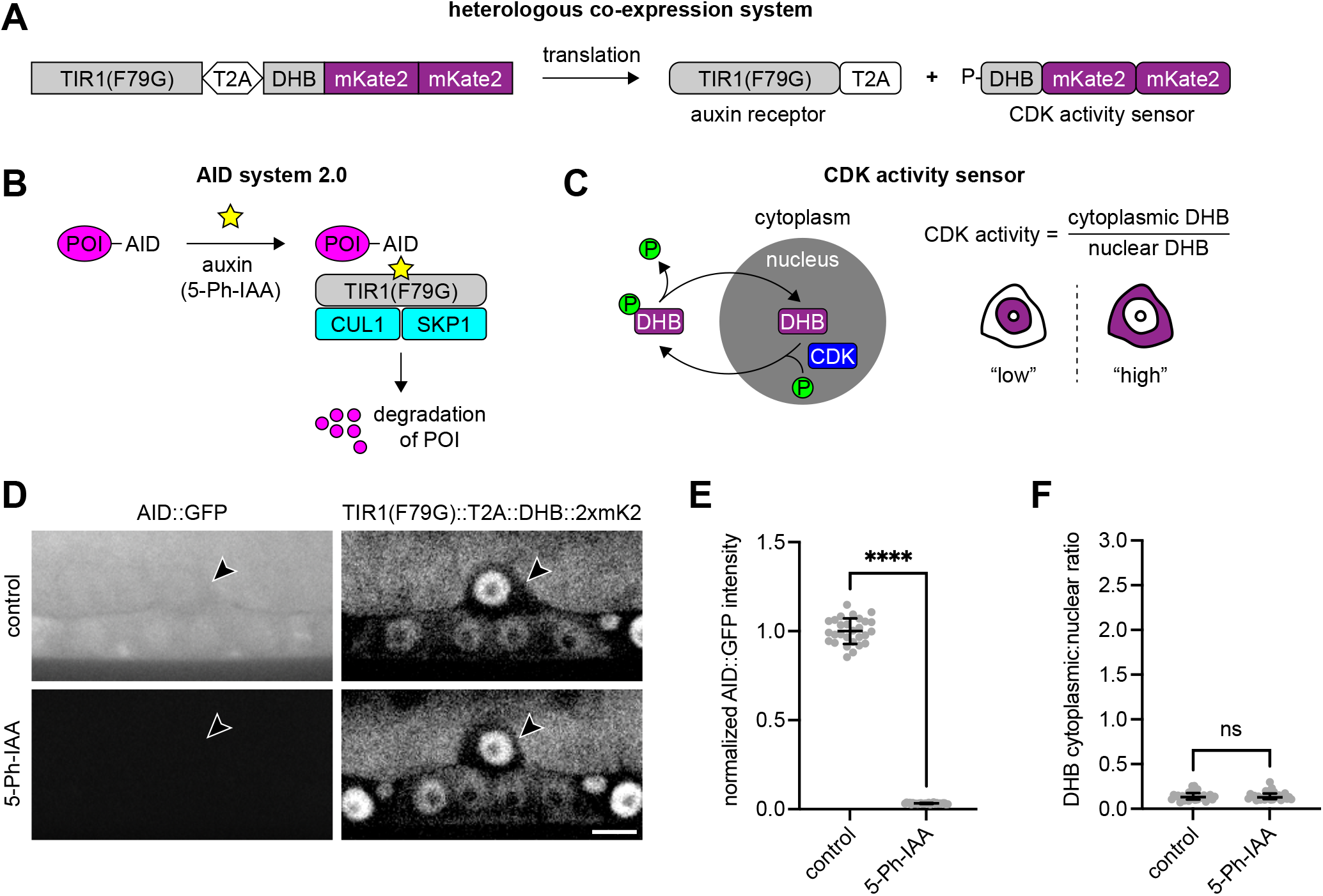
Conditional protein degradation and tracking of cell-cycle state in *C. elegans*. A. A bicistronic construct encoding TIR1(F79G) and DHB::2xmKate2 via a self-cleaving T2A peptide. B. The second version of the AID system requires a minimal AID tag on the protein of interest (POI), expression of the F-box mutant protein TIR1(F79G), and exogenous exposure to 5-Ph-IAA. When 5-Ph-IAA is present, TIR1(F79G) forms a functional E3 ligase complex with endogenous CUL1 and SKP1, which subsequently triggers the proteasomal degradation of the AID-tagged POI. C. The CDK activity sensor is a fragment of human DNA helicase B (DHB) fused to one or more fluorescent proteins. An increase in the cytoplasmic-to-nuclear ratio of fluorescent DHB is indicative of cell-cycle progression. In contrast, post-mitotic cells retain their nuclear DHB signal. D. Micrographs of mid-L3 larvae at the time of AC invasion expressing AID::GFP and TIR1(F79G)::T2A::DHB::2xmKate2 in the absence (top) and presence (bottom) of 5-Ph-IAA. Treatment was initiated at the L1 larval stage. E. Normalized AID::GFP intensity following 5-Ph-IAA treatment. Data presented as the mean with SD (n = 28 animals per treatment). P < 0.0001 as calculated by the Welch’s t test. F. Cytoplasmic-to-nuclear ratios of DHB::2xmKate2 following 5-Ph-IAA treatment. Data presented as the median with interquartile range (n = 28 animals per treatment). ns as calculated by the Mann-Whitney test.

## Results

### AIDHB: A heterologous co-expression system to degrade target proteins and monitor the cell cycle

The auxin-inducible degron (AID) system enables rapid degradation of *C. elegans* proteins (Ashley et al., 2021; Hills-Muckey et al., 2021; Martinez et al., 2020; Negishi et al., 2021; Sepers et al., 2022; Zhang et al., 2015). It requires a minimal AID tag on the protein of interest (POI), expression of the *Arabidopsis* F-box protein TIR1, and exogenous exposure to the plant hormone auxin. When auxin is present, TIR1 interacts with CUL1 and SKP1 to form an E3 ligase complex that ubiquitinates the AID-tagged POI for proteasomal degradation (Fig. 1B). Here, we used the second iteration of the AID system (Hills-Muckey et al., 2021; Negishi et al., 2021), which utilizes a TIR1(F79G) mutant protein and modified auxin (5-Ph-IAA), to limit leaky degradation (Martinez et al., 2020).

We co-expressed TIR1(F79G) with a small fragment of human DNA helicase B (DHB) fused to two copies of mKate2 (DHB::2xmKate2) (Fig. 1A). Co-expression was achieved using a single construct that contains the ubiquitous *rpl-28* promoter and a self-cleaving T2A peptide that separates both transgenes (Hills-Muckey et al., 2021). DHB::2xmKate2 serves as a CDK activity sensor for live-cell imaging (Adikes et al., 2020) (Fig. 1A,C). CDK activity is visualized by diffusion of fluorescent DHB into the cytoplasm from the nucleus, and it can be measured by quantifying the cytoplasmic-to-nuclear ratio of DHB signal (Fig. 1C). Because this ratio is used as a proxy for cell-cycle state, the combined AID and DHB system, which we refer to as AIDHB, allows us to degrade POIs and determine the effect on the cell cycle.

To test the AIDHB approach, animals with AID::GFP under the control of the ubiquitous *eft-3* promoter were given 5-Ph-IAA at the L1 larval stage. These animals were subsequently imaged and quantified at the mid-L3 (P6.p four-cell) larval stage when anchor cell (AC) invasion normally occurs (Fig. 1D). Control animals show high GFP abundance in the AC, whereas animals treated with auxin show a significant loss of AC GFP (Fig. 1E). Further, DHB localization in the AC appears to be unchanged between treatments and controls, i.e., in a CDK-low state (Fig. 1F). These data indicate that AIDHB can robustly degrade a functionally inert AID-tagged protein without affecting the cell cycle.

### Auxin-inducible degradation of EGL-43 prior to AC specification phenocopies *egl-43(RNAi)*

The null phenotype of *egl-43* includes embryonic lethality (Hwang et al., 2007) and L1 larval arrest (Rimann and Hajnal, 2007). RNAi directed against *egl-43* bypasses these phenotypes, which has revealed a role for EGL-43 in AC specification and invasion (Deng et al., 2020; Hwang et al., 2007; Matus et al., 2010; Medwig-Kinney et al., 2020; Rimann and Hajnal, 2007; Wang et al., 2014). Specifically, *egl-43(RNAi)* leads to the formation of two ACs and/or post-specification defects such as AC proliferation and failure to breach basement membrane (BM).

The conditionality of AIDHB should also allow us to avoid the developmental defects associated with *egl-43* null mutants. To explore this, we examined AC phenotypes using AIDHB with a new internally AID-tagged allele of *egl-43* that targets the long and short isoforms of endogenous EGL-43 (Fig. 2A), as these isoforms are thought to function redundantly (Medwig-Kinney et al., 2020). We also introduced endogenous alleles of *lag-2* (LAG-2::P2A::H2B::mTurquoise2) (Medwig-Kinney et al., 2022) and *lam-2* (LAM-2::mNeonGreen) (Jayadev et al., 2019) to label the AC and BM, respectively. Animals expressing all markers were treated with 5-Ph-IAA as L1 larvae and showed the proliferative AC phenotype (>2 ACs) in 24/32 animals (Fig. 2B-D). Of those animals, there was nearly an 88% defect in AC invasion. In 5/32 animals, two ACs formed without BM invasion. The two-AC phenotype is either due to a defect in specification, loss of the post-mitotic state, or both. Nonetheless, these data demonstrate that auxin-induced degradation of EGL-43 prior to AC specification resembles the AC phenotypes we and others have observed with *egl-43(RNAi)* (Deng et al., 2020; Medwig-Kinney et al., 2020).

**Figure 2.**
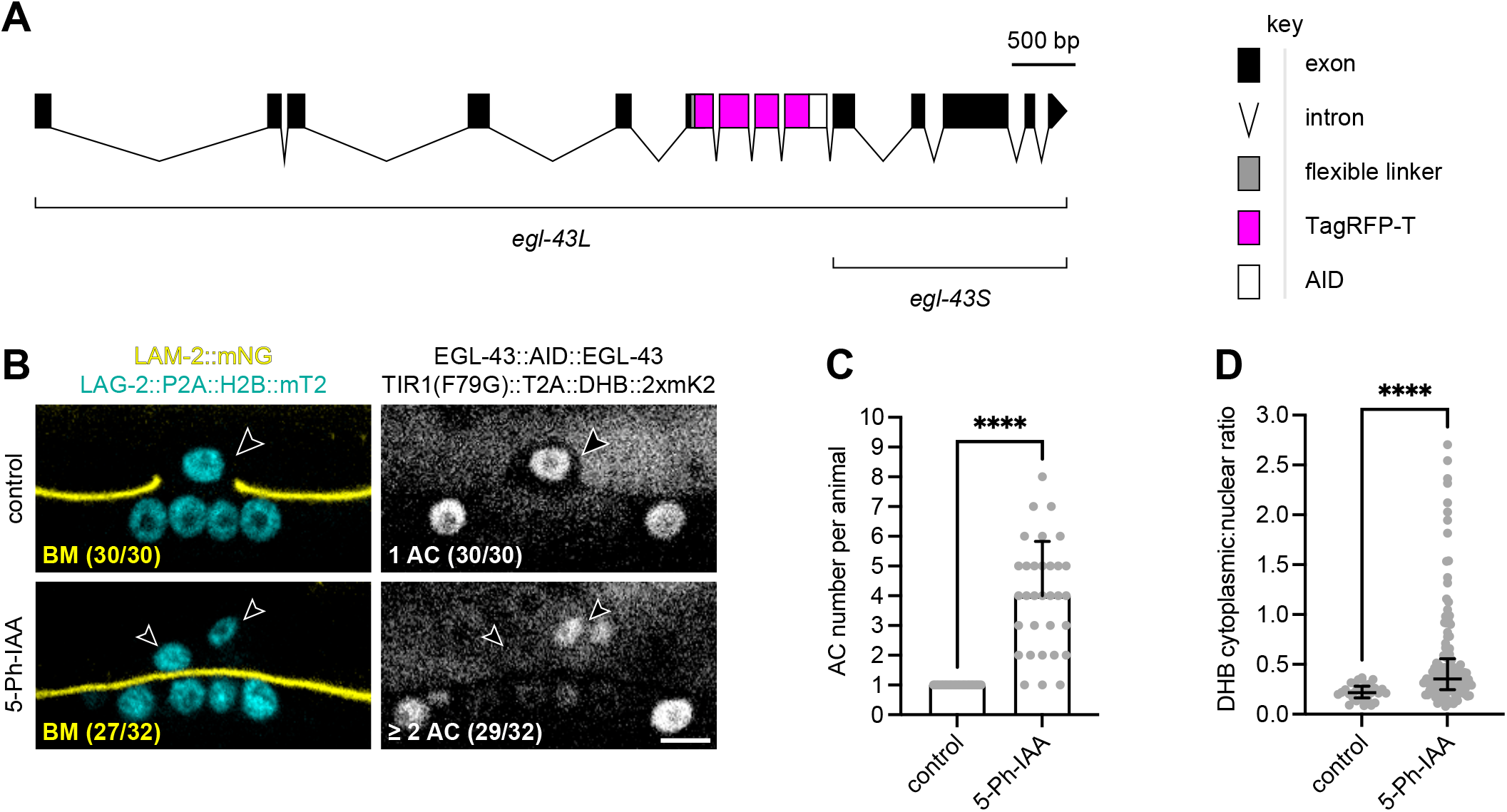
Robust degradation of EGL-43 produces the expected AC phenotypes. A. A schematic of the endogenously tagged AID allele of *egl-43*. This allele is hereafter referred to as EGL-43::AID::EGL-43, because TagRFP-T is undetectable above background levels of fluorescence. B. Micrographs of L3 larvae at the time of AC invasion expressing LAG-2::P2A::H2B::mTurquoise2 and LAM-2::mNeonGreen (left) as well as TIR1(F79G)::T2A::DHB::2xmKate2 and EGL-43::AID::EGL-43 (right) in the absence (top) and presence (bottom) of 5-Ph-IAA. Treatment was initiated at the L1 larval stage prior to AC specification, leading to defects in AC specification and AC invasion. C. Number of ACs per animal following 5-Ph-IAA treatment. Data presented as the mean with SD (n ≥ 30 animals per treatment). P < 0.0001 as calculated by the Welch’s t test. D. Cytoplasmic-to-nuclear ratios of DHB::2xmKate2 following 5-Ph-IAA treatment. Data presented as the median with interquartile range (n ≥ 30 animals per treatment). P < 0.0001 as calculated by the Mann-Whitney test.

### LIN-12 expression is not required for AC proliferation

AC specification is determined by a stochastic Notch signaling event between two equipotent cells (Greenwald et al., 1983). The cell that strongly expresses the transmembrane receptor LIN-12 becomes a ventral uterine cell, and the cell that strongly expresses its ligand, LAG-2, becomes the AC. In the absence of LIN-12, as in a *lin-12* null mutant, both cells become ACs. To further test AIDHB, we combined it with an endogenous allele of *lin-12* tagged at the C-terminus with mNeonGreen::AID (Pani et al., 2022). We also included LAG-2::P2A::H2B::mTurquoise2 as an AC marker. As expected (Deng et al., 2020), control animals showed no LIN-12 in the post-specified AC. Similar to the *lin-12* null mutant, auxin-induced degradation of LIN-12 in the L1 larval stage, prior to AC specification, resulted in the two-AC phenotype in 28/29 animals at the time of AC invasion (Fig. S1). Additionally, visualization of DHB in auxin-treated animals showed two post-mitotic ACs with low CDK activity, providing further evidence that loss of LIN-12 results in the generation of two ACs.

Recently, it was concluded that EGL-43 maintains the post-mitotic state of the AC by repressing LIN-12 (Deng et al., 2020). While LIN-12::GFP expression in proliferating ACs after *egl-43* or *nhr-67* RNAi was a striking result, only double RNAi directed against *egl-43* and *lin-12* suppressed the AC proliferation phenotype. Because the efficiency of double RNAi can be low (Min et al., 2010), we decided to pair AIDHB with RNAi. We exposed L1 larvae expressing AIDHB, LIN-12::mNeonGreen::AID, and LAG-2::P2A::H2B::mTurquoise2 to *egl-43(RNAi)* with and without 5-Ph-IAA. At the time of AC invasion, 30/30 auxin-treated animals and 26/30 control animals displayed the proliferative AC phenotype (Fig. 3A-C). In addition, the total number of ACs nearly doubled in auxin-treated animals compared to controls (n = 196 vs. 118). The higher total is expected for animals with two post-specified ACs that then entered the cell cycle and proliferated. Lastly, we confirmed the presence of LIN-12::mNeonGreen::AID in proliferating ACs of auxin controls after *egl-43(RNAi)* (Fig. S2), which localized to the cell membrane in 117/118 cases (see Discussion). Taken together, we conclude that LIN-12 is not required for AC proliferation.

**Figure 3.**
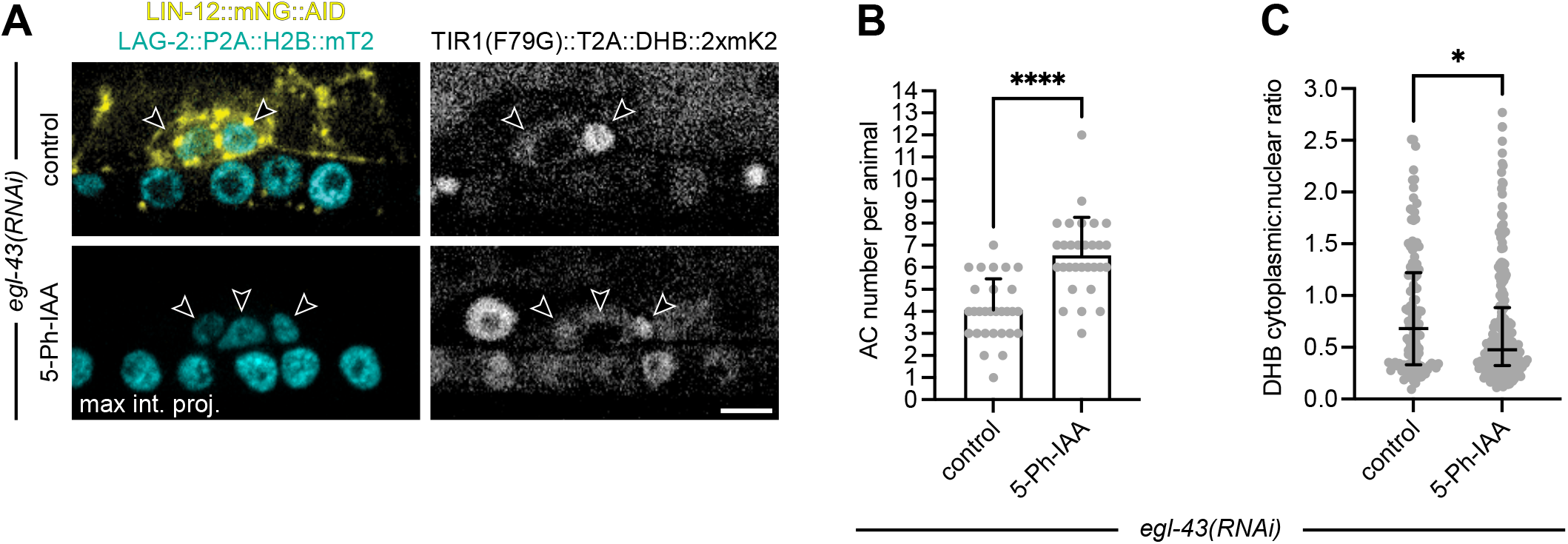
In the absence of EGL-43, LIN-12 is not required for AC proliferation. A. Micrographs of L3 larvae at the time of AC invasion expressing LAG-2::P2A::H2B::mTurquoise2 and LIN-12::mNeonGreen::AID (left) as well as TIR1(F79G)::T2A::DHB::2xmKate2 (right) after *egl-43(RNAi)* in the absence (top) and presence (bottom) of 5-Ph-IAA. All treatments were initiated at the L1 larval stage. B. Number of ACs per animal following *egl-43(RNAi)* and 5-Ph-IAA treatment. Data presented as the mean with SD (n ≥ 29 animals per treatment). P < 0.0001 as calculated by the Welch’s t test. C. Cytoplasmic-to-nuclear ratios of DHB::2xmKate2 following 5-Ph-IAA treatment. Data presented as the median with interquartile range (n ≥ 29 animals per treatment). P = 0.0378 as calculated by the Mann-Whitney test.

## Discussion

In this study, we built a tool called AIDHB to pair conditional protein degradation with visualization of cell-cycle state. We show that AIDHB can robustly degrade a non-functional AID::GFP protein without affecting the cell cycle of our cell of interest, the invasive AC. As a proof of concept, we targeted an AID-tagged allele of *egl-43* or *lin-12* for degradation beginning in the L1 larval stage before AC specification. These experiments produced the expected AC phenotypes observed with either LIN-12 or EGL-43 depletion. Whereas loss of LIN-12 results in the formation of two ACs due to a defect in AC specification (Greenwald et al., 1983), loss of EGL-43 leads to defects in AC specification and/or AC invasion (Deng et al., 2020; Hwang et al., 2007; Matus et al., 2010; Medwig-Kinney et al., 2020; Rimann and Hajnal, 2007; Wang et al., 2014). Finally, we sought to test the efficacy of combining AIDHB with RNAi, allowing us to reexamine the relationship between EGL-43 and LIN-12 during AC invasion. Recent work has shown that EGL-43 represses LIN-12 to maintain the post-mitotic state of the AC (Deng et al., 2020). Although we were able to confirm that *egl-43(RNAi)* results in ectopic *lin-12* expression in proliferating ACs, we did not observe localization in the nucleus. This suggests that ectopic LIN-12 may not be representative of active Notch signaling (Medwig-Kinney et al., 2022; Pani et al., 2022). When we combined AIDHB and RNAi to deplete LIN-12 and EGL-43, respectively, we found that EGL-43-deficient ACs were able to proliferate in the absence of LIN-12. This is in contrast to animals treated with double RNAi directed against *egl-43* and *lin-12* (Deng et al., 2020), but the efficiency of RNAi can suffer when more than one gene is targeted (Min et al., 2010). Together, our results reveal that LIN-12 is not required for AC proliferation.

What promotes AC proliferation following loss of EGL-43, HLH-2, or NHR-67 remains an open question. Interestingly, in the presence of EGL-43, AC-specific expression of the Notch intracellular domain (NICD) can force the AC to proliferate (Deng et al., 2020). The NICD is the functionally active component of LIN-12 that is released into the nucleus after a series of proteolytic cleavages (Falo-Sanjuan and Bray, 2020). It should be noted, however, that NICD-driven AC proliferation may require a deletion of the NICD C-terminal PEST domain (Nusser-Stein et al., 2012). NICD constructs lacking this domain are potentially resistant to endogenous mechanisms of degradation. Thus, our findings, coupled with these observations, suggests that AC proliferation in this context is a neomorphic phenotype. This is consistent with other cases where ectopic NICD expression can induce proliferation (Kwon et al., 2014; Kwon et al., 2016; Valdez et al., 2012). Based on ChIP-seq data, there are putative EGL-43 binding sites in the *lin-12* locus (Deng et al., 2020). The emergence of CRISPR/Cas9 as a gene-editing tool in *C. elegans* (Vicencio and Cerón, 2021) should facilitate the modification of these binding sites, helping to elucidate the relationship between EGL-43 and LIN-12 during AC invasion.

In summary, we (i) created a heterologous co-expression system called AIDHB, which we paired with RNAi, (ii) generated a new AID-tagged allele of *egl-43*, and (iii) postulate that in the absence of EGL-43, LIN-12 expression is not necessary for AC proliferation. It is our hope that investigators will utilize AIDHB to interrogate the function of diverse proteins that may be required for cell-cycle driven cellular behaviors.

## Materials and Methods

### Strains

Strains were maintained under standard culture conditions (Brenner, 1974). The following alleles were used in this study: LG I: *bmd284[rpl-28p::TIR1(F79G)::T2A::DHB::2xmKate2]*; LG II: *wy1514[egl-43::TagRFP-T::AID::egl-43]*; LG III: *ljf33[lin-12::mNeonGreen::AID]* (Pani et al., 2022); LG IV: *ieSi58[eft-3p::AID::GFP]* (Zhang et al., 2015); LG V: *bmd202[lag-2::P2A::H2B::mTurquoise2]* (Medwig-Kinney et al., 2022), *bmd299[lag-2::P2A::H2B::mTurquoise2]*; LG X: *qy20[lam-2::mNeonGreen]* (Jayadev et al., 2019).

### Generation of the transgenic bmd284 allele

To clone pWZ259 (rpl-28p::TIR1(F79G)::T2A::DHB::2xmKate2), pWZ192 (NotI-ccdB-SphI-DHB::2xmKate2) was double digested with NotI and SphI to excise ccdB and a PCR product representing rpl-28p::TIR1(F79G)::T2A was amplified from plasmid pCMH2123 using primers DQM1136 and DQM1137. pWZ259 was constructed by Gibson assembly (NEB) using the backbone from pWZ192 and the PCR product from pCMH2123. After sequence confirmation, pWZ259 was used as a repair template for insertion into the genome at a safe harbor site on chromosome I corresponding to the MosSCI insertion site ttTi4348 (Frøkjær-Jensen et al., 2012). pAP082 was used as the sgRNA plasmid for chromosome I insertion via CRISPR/Cas9 (Pani and Goldstein, 2018). Young adults were transformed using standard microinjection techniques and integrants were identified through the SEC method (Dickinson et al., 2015).

### Generation of the endogenous wy1514 allele

A repair template containing TagRFP-T::AID with homology at the 5’ and 3’ ends to the *egl-43* locus was PCR amplified and purified using a PCR purification kit (Qiagen). 3 μl of 10 μM tracRNA (IDT) was incubated with 0.5 μl of 100 μM of a crRNA (IDT) targeting exon 6 of the *egl-43* locus at 95°C for 5 minutes, followed by 25°C for 5 minutes. Following incubation, the mixture was incubated with 0.5 μl of Cas9 protein (IDT) at 37°C for 10 minutes. Repair template and a co-injection marker (pRF4) were added to the mixture to a final concentration of 200 ng/μl and 50 ng/μl, respectively. Young adult worms were transformed using standard microinjection techniques and progeny were genotyped for successful insertions (Paix et al., 2015).

### Auxin treatment

Synchronized L1 larvae were plated on NGM plates containing 0.1 mM 5-Ph-IAA (MCE) and fed either OP50 or *egl-43(RNAi)*. The *egl-43(RNAi)* feeding construct was published previously (Medwig-Kinney et al., 2020), and it silences the expression of both the long and short isoform of EGL-43. 0.1% ethanol was used as an auxin control. All animals were analyzed at the mid-L3 (P6.p four-cell) larval stage when AC invasion occurs.

### Image acquisition

Images were collected using a custom-built spinning disk confocal microscope (Nobska Imaging), which was configured for automation with Metamorph software (Molecular Devices). This confocal consists of a Hamamatsu ORCA EM-CCD camera mounted on an upright Zeiss Axio Imager.A2 with a Borealis-modified Yokogawa CSU-10 spinning disk scanning unit and a Zeiss Plan-Apochromat 100x/1.4 oil DIC objective. Animals were anesthetized for imaging by picking them into a drop of M9 on a 5% agarose pad containing 7 mM sodium azide and secured with a coverslip.

### Image processing and analysis

Acquired images were processed using ImageJ/Fiji (Schneider et al., 2012). AID::GFP fluorescence was quantified as previously described (Martinez and Matus, 2020). DHB::2xmKate2 ratios were quantified as previously described (Adikes et al., 2020). AC number was determined by counting AC nuclei (LAG-2::P2A::H2B::mTurquoise2). AC invasion was defined as the complete loss of BM (LAM-2::mNeonGreen) under the AC. Plots were generated using Prism software. Figures, and the cartoons within, were created using a combination of Adobe Photoshop and Illustrator.

## Acknowledgments

We thank Taylor Medwig-Kinney for generating *bmd298*, which is the precursor allele to *bmd299*, as previously described (Medwig-Kinney et al., 2022).

## Author Contributions

Conceptualization: M.A.Q.M., D.Q.M.; Methodology: M.A.Q.M., D.Q.M.; Formal Analysis: M.A.Q.M., A.A.M., C.Z.; Investigation: M.A.Q.M., A.A.M.; Resources: C.Y., W.Z., K.S.; Writing – original draft preparation: M.A.Q.M., A.A.M.; Writing – review and editing: M.A.Q.M., D.Q.M.; Visualization: M.A.Q.M.

