## Supplemental Files for "A reevaluation of the relationship between EGL-43 (EVI1/MECOM) and LIN-12 (Notch) during *C. elegans* anchor cell invasion"

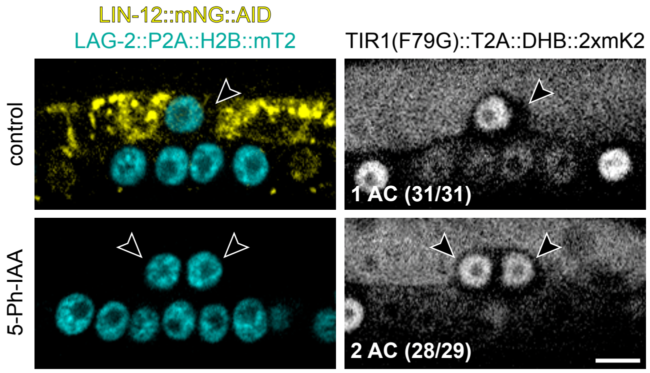
LIN-12::mNG::AID
LAG-2::P2A::H2B::mT2
TIR1(F79G)::T2A::DHB::2xmK2
control
5-Ph-IAA
1 AC (31/31)
2 AC (28/29)

**Figure S1. Robust degradation of LIN-12 produces the expected AC phenotype.**

Micrographs of mid-L3 larvae at the time of AC invasion expressing LAG-2::P2A::H2B::mTurquoise2 and LIN-12::mNeonGreen::AID (left) as well as TIR1(F79G)::T2A::DHB::2xmKate2 (right) in the absence (top) and presence (bottom) of 5-Ph-IAA. Treatment was initiated at the L1 larval stage prior to AC specification, resulting in the two-AC phenotype.

LIN-12::mNG::AID

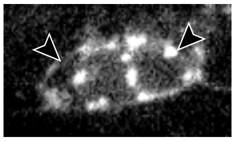

auxin control

*egl-43(RNAi)*

**Figure S2. EGL-43 depletion leads to ectopic LIN-12::mNG::AID expression on the AC membrane.**

A micrograph of a mid-L3 at the time of AC invasion expressing LIN-12::mNeonGreen::AID after *egl-43(RNAi)* in the absence of 5-Ph-IAA. Each treatment was initiated at the L1 larval stage.

**Table S1: Strains**

| Name | Genotype | Source |
|---|---|---|
| DQM1125 | *bmd284 I*; *ljf33 III*; *bmd299 V* | This paper |
| DQM1159 | *bmd284 I*; *ieSi58 IV* | This paper |
| DQM1233 | *bmd284 I*; *wy1514 II*; *bmd202 V*; *qy20 X* | This paper |

**Table S2: Primers**

| Name | Sequence (5' – 3') | Type | Amplicon | Template |
|---|---|---|---|---|
| DQM1136 | tgtaaaacgacggccagtgcggccgcGT TTGTGCAACAAATTGAGGAAG | Forward | rpl-28p::TIR1(F79G)::T2A | pCMH2123 |
| DQM1137 | caggtgacgtcgttggtcatgggcccTCC TGGGCCAGGATTCTC | Reverse | rpl-28p::TIR1(F79G)::T2A | pCMH2123 |
| CY419 | caataagacacgcgcgccgcccatccctc gtgcatcttccctttggggtgtgtcacaggg ctcattcttgtgacgtgcgcttcccaccttt aAcagtgtgttatcaatctccgtcttattca ttagtgaataaatattccaggacggttgga cgtgcccaacgtgtcaaagtcagGGAG CATCGGGAGCCTCAGGAGCat | Forward | egl-43::TagRFP-T::AID::egl-43 | pWZ203 |
| CY420 | agttttcaaaataataactaacctgtgttg ccatttgaagtatatgtggccaatatggca cggaacctaatccacctgctccgctcaatc cggcaagttgcgccatcaatggtttgtaga gtgcagtcatctcgcaaactggtcgatgct tcgttagtgccgcttgtgatggcatGGCT CCGCTAGCTCCTGACTTGACG | Reverse | egl-43::TagRFP-T::AID::egl-43 | pWZ203 |

**Table S3: Plasmids**

| Name | Backbone | Description |
|---|---|---|
| pWZ259 | pWZ192 | rpl-28p::TIR1(F79G)::T2A::DHB::2xmKate2 |

**Table S4: Guides**

| Locus | Sequence (5' – 3') | Description |
|---|---|---|
| *egl-43* | GACGGAGATTGATAACACAC | Located upstream of exon 6 |
